# Selective linkage of mitochondrial enzymes to intracellular calcium stores differs between human induced pluripotent stem cells, neural stem cells and neurons

**DOI:** 10.1101/2020.06.20.162040

**Authors:** Huanlian Chen, Ankita Thakkar, Abigail C. Cross, Hui Xu, Aiqun Li, Dan Pauli, Scott A. Noggle, Laken Kruger, Travis T. Denton, Gary E. Gibson

**Affiliations:** Burke Neurological Institute, Brain and Mind Research Institute, Weill Cornell Medicine, White Plains, NY; The New York Stem Cell Foundation Research Institute, New York, NY; Department of Pharmaceutical Sciences, Washington State University, College of Pharmacy and Pharmaceutical Sciences, Spokane, WA; Department of Genetics and Genomic Sciences, Icahn School of Medicine at Mount Sinai, One Gustave L. Levy Place, New York, NY

**Keywords:** Alzheimer’s disease, mitochondria, endoplasmic reticulum, development, stem cells, alpha-ketoglutarate dehydrogenase complex, pyruvate dehydrogenase complex, calcium stores, tricarboxylic acid cycle

## Abstract

The coupling of the endoplasmic reticulum (ER) with mitochondria modulates neuronal calcium signaling. Whether this link changes with neuronal development is unknown. The current study first determined whether ER calcium stores are similar during development of human neurons, and then tested if the ER/mitochondrial coupling varied with development. The release of ER calcium to the cytosol by the IP_3_ agonist bradykinin was determined in human induced-pluripotent stem cells (iPSC), neural stem cells (NSC) and neurons. The concentration dependence for the release of ER calcium was similar at different stages of development. Metabolism changes dramatically with development. Glycolysis is the main energy source in iPSC and NSC whereas mitochondrial metabolism is more prominent in neurons. To test whether the coupling of mitochondria and ER changed with development, bombesin or bradykinin releasable calcium stores (BRCS) were monitored after inhibiting either of two key mitochondrial enzyme complexes: the alpha-ketoglutarate dehydrogenase complex (KGDHC) or the pyruvate dehydrogenase complex (PDHC). Inhibition of KGDHC did not alter BRCS in either iPSC or NSC. Inhibition of PDHC in neurons diminished BRCS whereas decreased KGDHC activity exaggerated BRCS. The latter finding may help understand the pathology of Alzheimer’s disease (AD). BRCS is exaggerated in cells from AD patients and KGDHC is reduced in brains of patients with AD. In summary, a prominent ER/mitochondrial link in neurons is associated with selective mitochondrial enzymes. The ER/mitochondrial link changes with human neuronal development and plausibly links ER calcium changes to AD.

## Introduction

The physiological and morphological connections between mitochondria and the endoplasmic reticulum (ER) are critical in the regulation of neuronal calcium signaling (Csordás *et al*. 1999; Hajnóczky *et al*. 1994; Rizzuto *et al*. 1993; Rizzuto *et al*. 1998; Karagas & Venkatachalam 2019) and in Alzheimer’s disease (AD). The IP_3_ receptors (IP_3_R) on the ER appear to be in close proximity to the mitochondria (Seitaj *et al*. 2018). The relative reliance on mitochondria and glycolysis varies in human induced pluripotent stem cells (iPSC), neural stem cells (NSC) and neurons. iPSC and NSC are very glycolytic, whereas metabolism in neurons is more mitochondrial (Intlekofer & Finley 2019; Khacho *et al*. 2019). However, little is known about calcium signaling during various stages of development of human neurons. Thus, these experiments test whether cytosolic calcium in human iPSC, NSC or neurons increases similarly in response to potassium depolarization or to IP_3_ agonists such as bradykinin (BRCS).

Considerable data suggest that releasable ER calcium stores are abnormal in AD. The calcium stores that are released by agonists that increase IP_3_ formation, such as bradykinin or bombesin are exaggerated in cells from AD patients, as well as in cell and animal models of AD. Bombesin or bradykinin releasable calcium stores (BRCS) are exaggerated in fibroblasts from patients with both genetic and non-genetic forms of AD (Ito *et al*. 1994), cultured fibroblasts from presenilin-1 (PS-1) knock-in mice (Leissring *et al*. 2000) and cultured neurons from 3×Tg AD mice (Stutzmann *et al*. 2004; Stutzmann *et al*. 2006). The abnormal proteins resulting from presenilin mutations interact with the IP_3_ receptor Ca^2+^ release channel and alter its stimulatory effect in response to IP_3_ (Cheung *et al*. 2008; Cheung *et al*. 2010). These calcium changes are likely pathologically important, therefore, modulating Ca^2+^ release is a reasonable therapeutic target (Schrank *et al*. 2019).

The activities of key mitochondrial enzymes decline in AD and these abnormalities may modulate the mitochondrial/ER interactions that accompany AD. In non-human tissues, inhibition of one of these enzymes, the alpha-ketoglutarate dehydrogenase complex (KGDHC) exaggerates BRCS in an AD-like manner. The precise mechanism is unknown, but reductions in KGDHC diminish GTP (Kiss *et al*. 2013), and GTP is known to modulate ER/mitochondrial interactions (Hajnóczky *et al*. 1994). Furthermore, diminished KGDHC causes release of mitochondrial proteins (Banerjee *et al*. 2016) and post-translational modification of non-mitochondrial proteins (Yang *et al*. 2019). The activity of the pyruvate dehydrogenase enzyme complex (PDHC), which links glycolysis and mitochondrial metabolism, also declines with AD, but the relation to the ER/mitochondrial interactions has never been examined.

Thus, the current experiments test the interactions of KGDHC with BRCS in human neurons at different stages of development and, additionally, compare the consequences of diminished KGDHC and PDHC activities in neurons. The results reveal that the consequences differ between the two enzymes and between human iPSC, NSC and neurons. The results support the hypothesis that treating the mitochondrial deficit related to AD may ameliorate calcium abnormalities in AD.

## Results

### Generation and characterization of iPSC, NSC and iPSC-derived neurons

All of the cell types were confirmed with their cell-specific markers (iPSCs: OCT4 and SSEA4; NSC: Nestin, SOX2, SOX1 and PAX6, iPSC derived neurons: MAP2, neurofilament and synaptophysin) (Figure-1). The identity of the iPSC-derived neurons was confirmed by depolarization of voltage gated calcium channels by 50 mM KCl. K^+^ depolarization increased intracellular calcium response in iPSC-derived neurons, but not in either iPSC or NSC (Figure 2). Thus, both the antibody marker studies and the depolarization indicated that our robust neuronal differentiation protocol yielded highly homogenous neurons.

**Figure 1.**
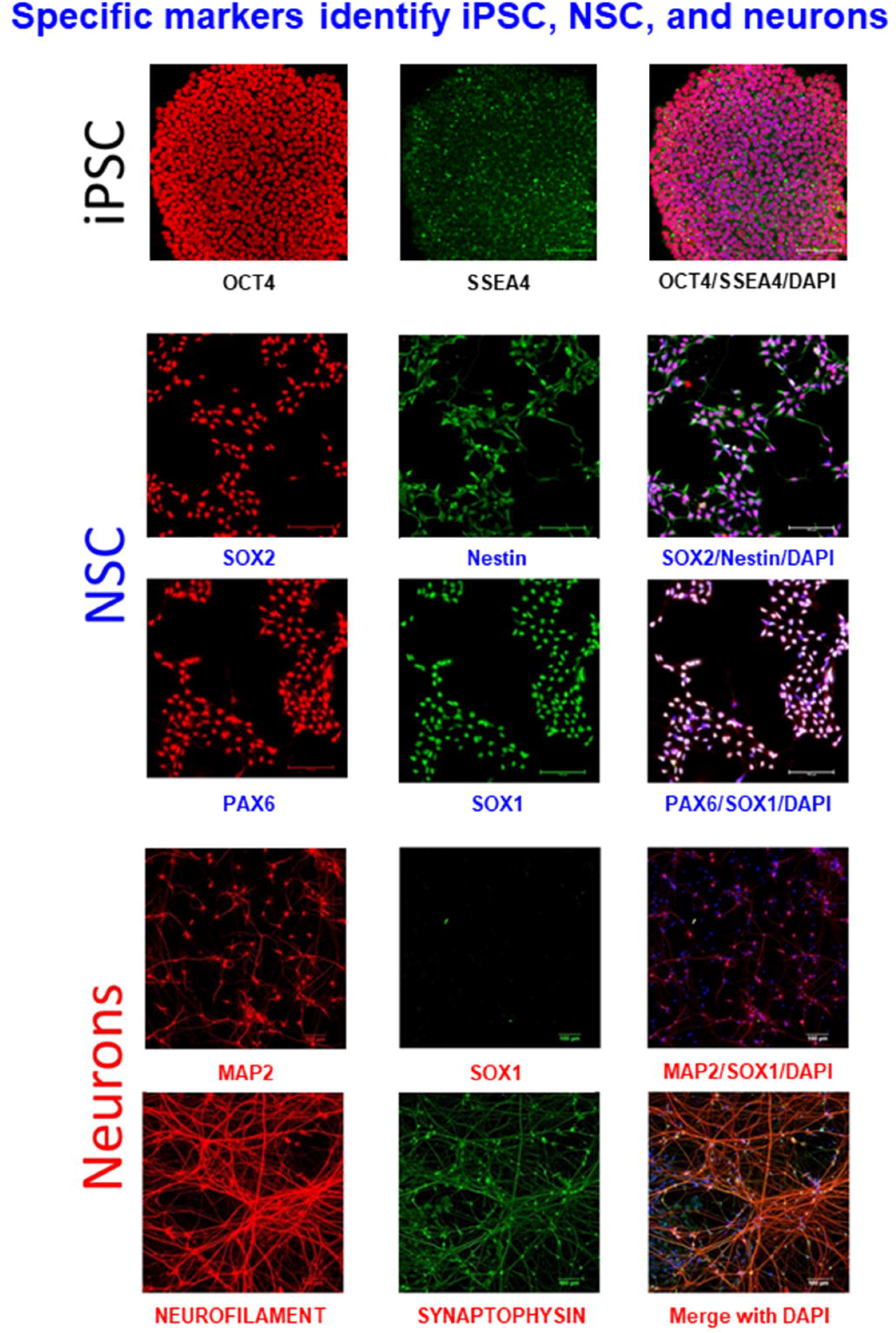
Immunocytochemical characterization of iPSC, NSC and iPSC derived neurons. NSC were differentiated from iPSC (passage 30-35) after 7 days in culture. NSC were cultured for five passages before initiating neuronal differentiation. Neurons. Cells were differentiated for four weeks. Cell specific markers were confirmed in cells at the three stages of differentiation. Cells were cultured on Delta T dishes at different seeding densities and culture times (iPSC, 2 × 10^5^ cell/dish for two days; NSC, 2 × 10^5^ cell/dish for two days; neurons, 1 × 10^5^ cell/dish for four weeks). Markers were tested by immunocytochemistry (iPSCs: OCT4 and SSEA4; NSC: Nestin, SOX2, SOX1 and Pax6, iPSC derived neurons: MAP2, neurofilament and synaptophysin).

**Figure 2.**
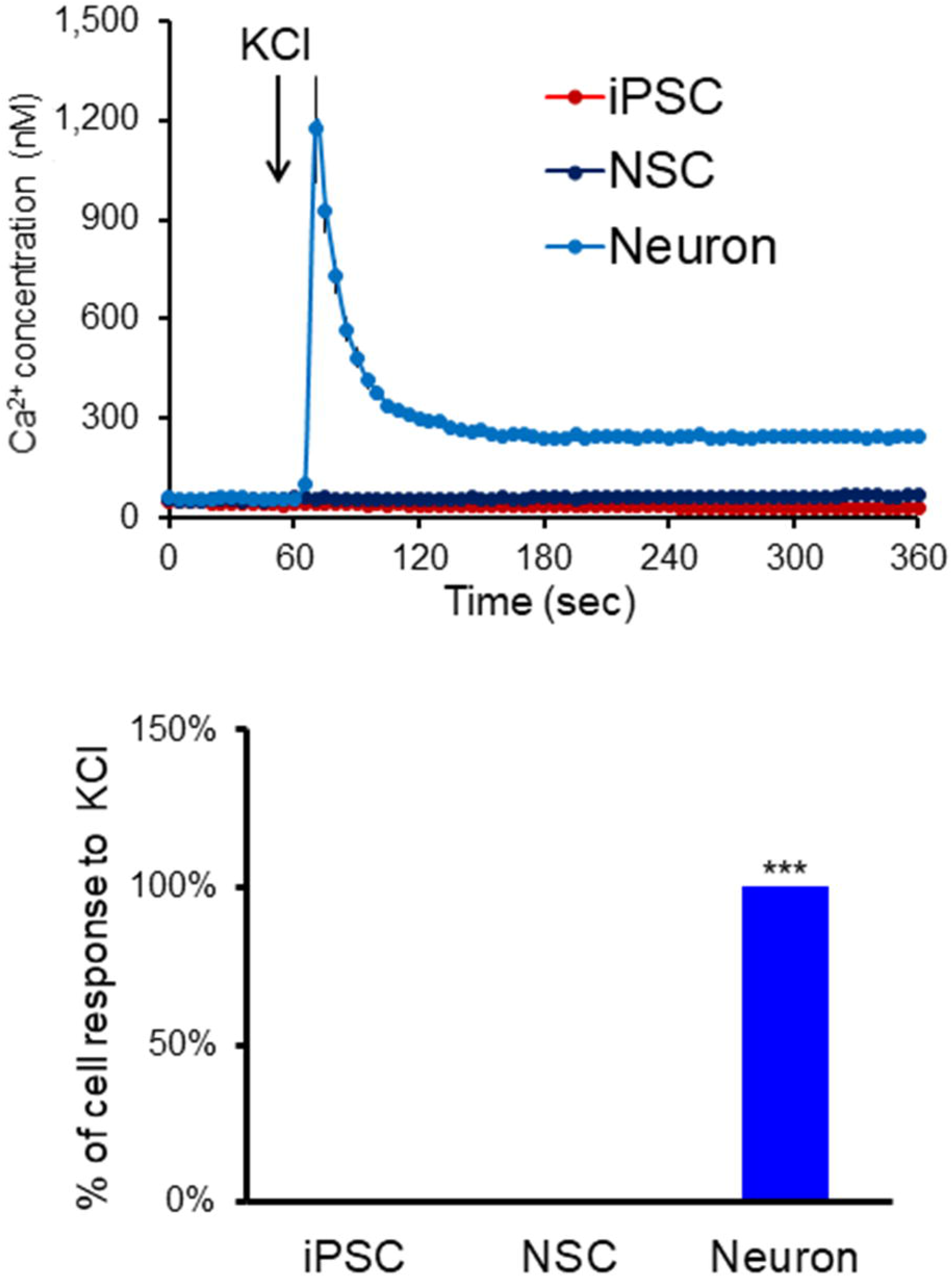
Characterization of voltage gated calcium channels in iPSC-derived cells. Functional characterization of the neurons was done by assessing the responses of [Ca^2+^]_i_ to depolarizing concentrations of K^+^ (KCl, 50 mM) in iPSC, NSC and iPSC derived neurons. Cells were loaded with Fura-2 AM (2 μM) in BSS for 60 min. After loading, the cells were rinsed with BSS and [Ca^2+^]_i_ was measured for one min of basal response and for another 5 min after adding KCl (50 mM): (A) The temporal response of [Ca^2+^]_i_ after adding KCl at 1 min. (B) the percentage of cell response to KCl in the three types of cells. Values are means ± SEM (n=106, 92 and 72 cells in iPSC, NSC, and iPSC-derived neurons, respectively). The error bars for the iPSC and NSC were too small to show. *** indicates values vary significantly (p<0.05) from the other groups by ANOVA followed by Student Newman Keul’s test.

### BRCS in iPSC, NSC and iPSC-derived neurons

The dose response to bradykinin determined whether BRCS varied with differentiation. Cells at each stage of differentiation had a dose dependent response to bradykinin. The patterns of change in iPSC (Figure 3) and neurons (Figure 5) were very similar. The [Ca^2+^]_i_ peaked and then went back to the basal [Ca^2+^]_i_ within two minutes after bradykinin addiction. However, the NSC responded differently than the other cells. The [Ca^2+^]_i_ did not return to basal [Ca^2+^]_i_ even after 5 min of bradykinin addition (Figure 4).

**Figure 3.**
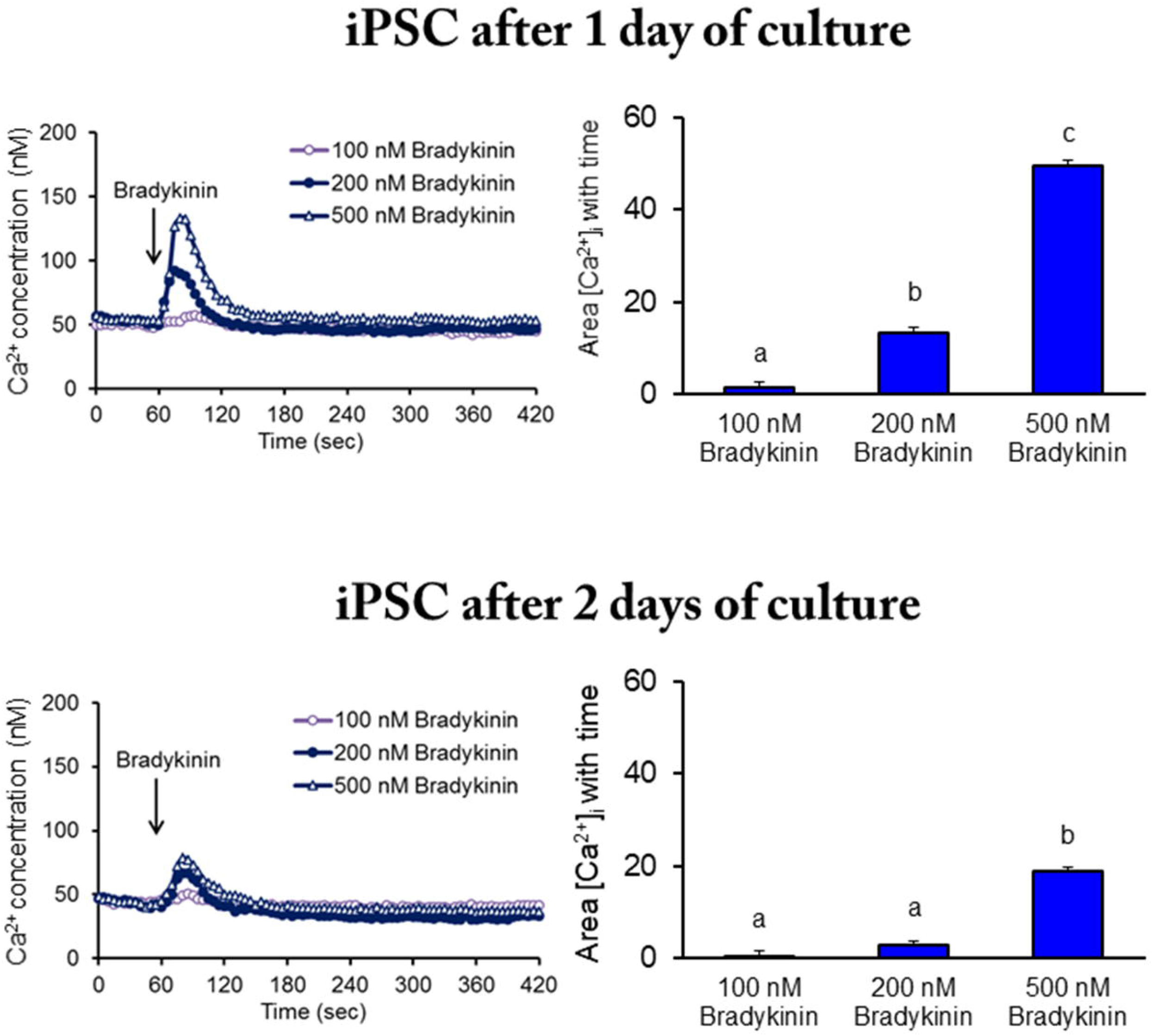
BRCS in iPSC. iPSC were cultured on Delta T dishes for one or two days. The cells were loaded with Fura-2 AM (2 μM) in BSS for 60 min for dye loading. The media were changed to calcium free BSS and the calcium measurements were initiated. Bradykinin (100, 200 or 500 nM) was added after 1 min of basal [Ca^2+^]_i_ measurement. The left panel shows the tracings and the right panel shows the integration of the [Ca^2+^]_i_ peak over the 5 min interval after bradykinin addition. Values are means ± SEM (n=355 and 250 cells in one and two days of culture, respectively). Different letters indicate values vary significantly (p<0.05) from the other groups by ANOVA followed by Student Newman Keul’s test.

**Figure 4.**
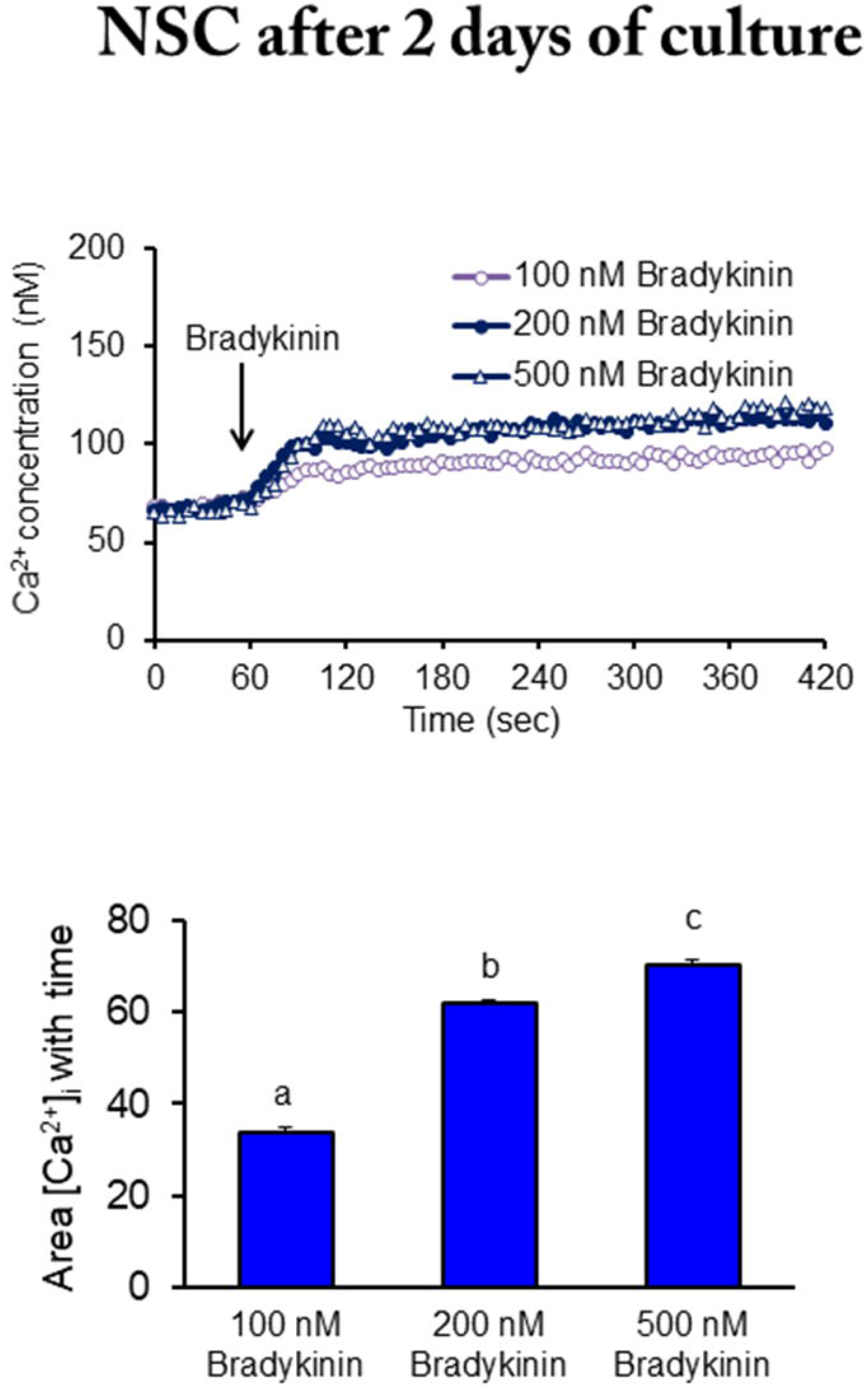
BRCS in NSC. NSC (passage 5) were cultured on Delta T dishes for two days. After two days of culture, the cells were loaded with Fura-2 AM (2 μM) in BSS for 60 min (Gibson *et al*. 2002). After loading, the media were changed to calcium free BSS. The calcium measurements were initiated and bradykinin (100, 200 or 500 nM) was added after 1 min of basal [Ca^2+^]_i_ measurement. The top panel shows the tracings and the bottom panel shows the integration of the [Ca^2+^]_i_ peak over the 5 min interval after bradykinin addition. Values are means ± SEM (n=490 cells). Different letters indicate values vary significantly (p<0.05) from the other groups by ANOVA followed by Student Newman Keul’s test.

**Figure 5.**
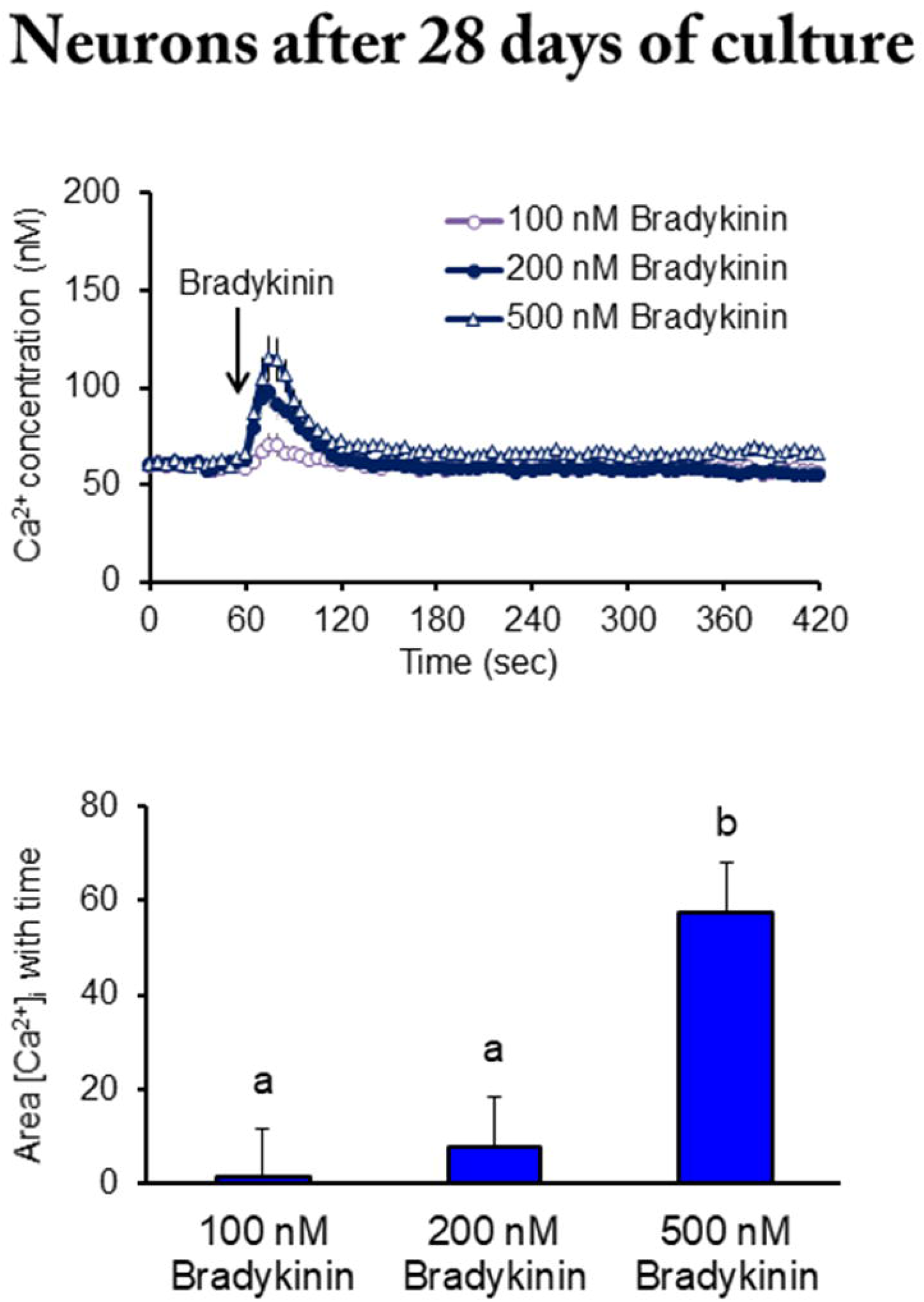
BRCS in iPSC-derived neurons. NSC (passage 5) were cultured on Delta T dishes. After four weeks of culture, the cells were loaded with Fura-2 AM (2 μM) in BSS for 60 min. After loading, the media were changed to calcium free BSS, and the calcium measurements were initiated. Bradykinin (100, 200 or 500 nM) was added after 1 min of basal [Ca^2+^]_i_ measurement. The top panel shows the tracings taken and the bottom panel shows the integration of the [Ca^2+^]_i_ peak over the 5 min interval after bradykinin addition. Values are means ± SEM (n=435 cells). Different letters indicate values vary significantly (p<0.05) from the other groups by ANOVA followed by Student Newman Keul’s test.

### The consequences of reducing KGDHC on BRCS in iPSC, NSC and iPSC-derived neurons

Cells were treated with the specific KGDHC inhibitor, trisodium succinyl phosphonate (SP, 500 μM), before adding bradykinin. First, the inhibitory effect of SP on KGDHC was confirmed with a histochemistry assay. In iPSC, 500 μM SP inhibited KGDHC activity by 67% after a one-hour treatment and by 75% after 24 hours of treatment (Figure 6). In NSC, SP inhibited KGDHC activity by 89% after one hour of treatment and by 94% after 24 hours of treatment (Figure 7). In iPSC-derived neurons, 500 μM SP inhibited KGDHC activity by 87% after one hour of treatment and by 83% after 24 hours of treatment (Figure 8). Reducing KGDHC with SP for one hour and 24 hours did not alter BRCS in iPSC (Figure 6) nor NSC (Figure 7). However, reducing KGDHC in iPSC-derived neurons exaggerated BRCS by 69% and 144% after a one-hour or 24-hour treatment, respectively (Figure 8).

**Figure 6.**
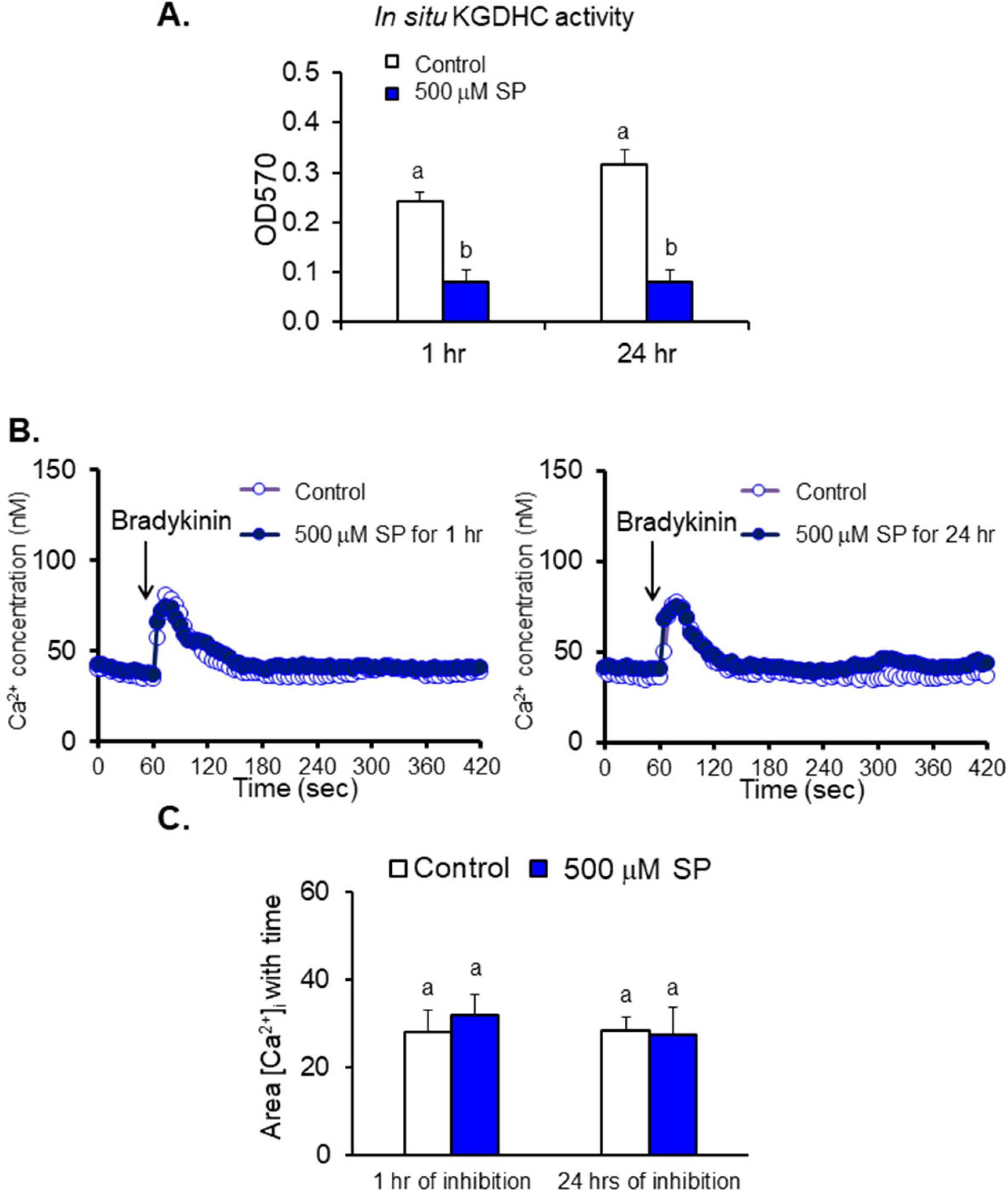
Inhibiting KGDHC did not alter BRCS in iPSC. iPSC were cultured in 24-wells plate at a seeding density of 1 × 10^5^ cells/well for two days. After two days of culture, cellular KGDHC activity was measured by histochemistry (Panel A). iPSC were cultured on Delta T dishes for one or two days. After culture, the cells were pre-treated with trisodium succinyl phosphonate (SP, 500 μM) for one hour (in BSS) or 24 hours (in completed media). After 1 hour of Fura-2 loading, the media were changed to calcium free BSS, and the calcium measurements were initiated and bradykinin (500 nM) was added after 1 min of basal [Ca^2+^]_i_ measurement. (Panel B). Values are means ± SEM (n=261 and 272 cells for one hour and 24 hours of treatment, respectively). Different letters indicate values vary significantly (p<0.05) from the other groups by ANOVA followed by Student Newman Keul’s test (Panel C).

**Figure 7.**
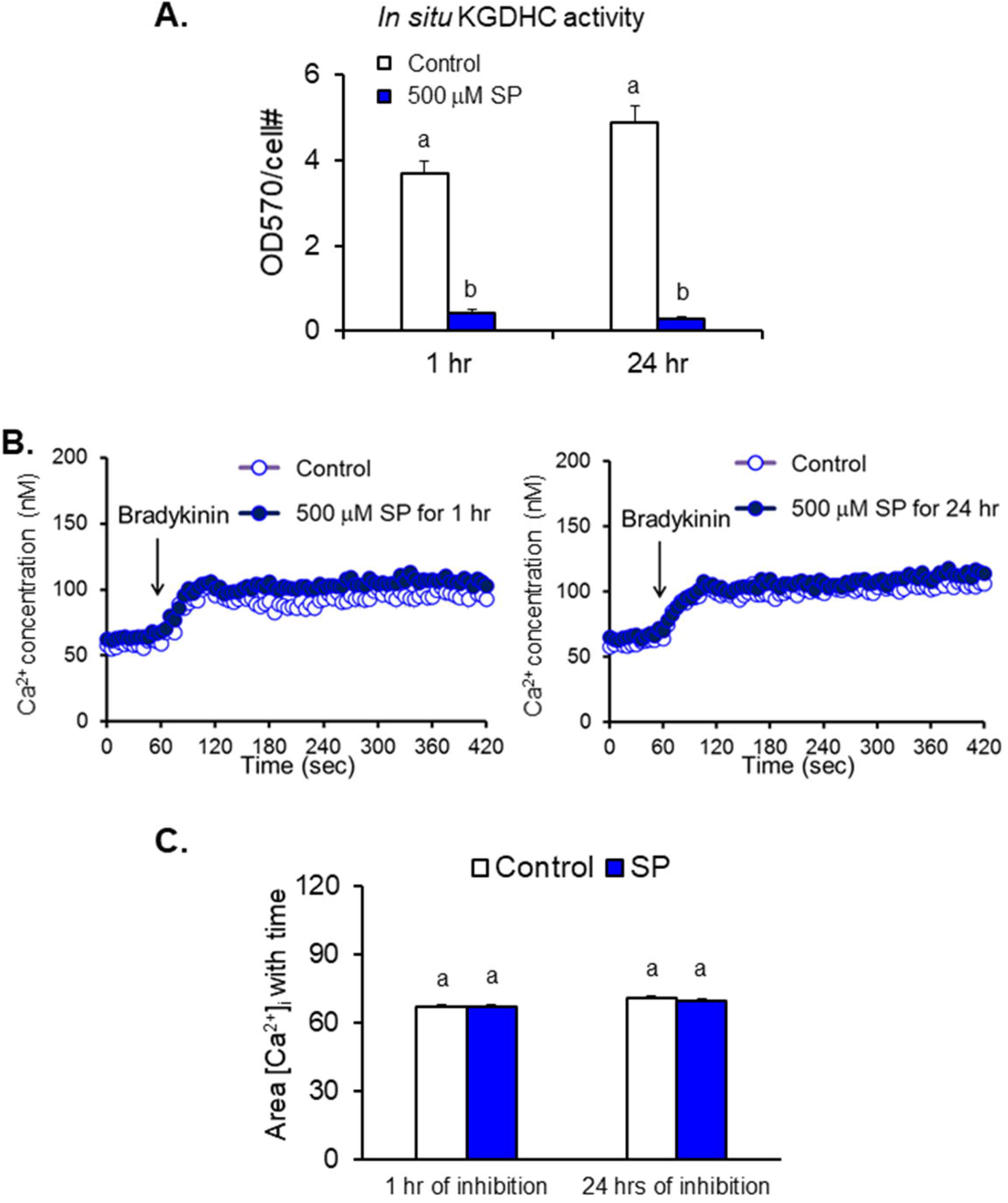
Reducing KGDHC did not alter BRCS in NSC. NSC were cultured on 24-well plate at the seeding density of 1 × 10^5^ cell/well for two days. After two days of culture, KGDHC activity was measured by histochemistry (Panel A). NSC were cultured on Delta T dishes for two days. After culture, the cells were pre-treated with trisodium succinyl phosphonate (SP, 500 μM) one hour (in BSS) or for 24 hours (in completed media). After 1 hour of Fura-2 loading, the media were changed to calcium free BSS, and the calcium measurements were initiated and bradykinin (500 nM) was added after 1 min of basal [Ca^2+^]_i_ measurement (Panel B). Values are means ± SEM (n=208 and 205 cells for one hour and 24 hours of treatment, respectively). Different letters indicate values vary significantly (p<0.05) from the other groups by ANOVA followed by Student Newman Keul’s test (Panel C).

**Figure 8.**
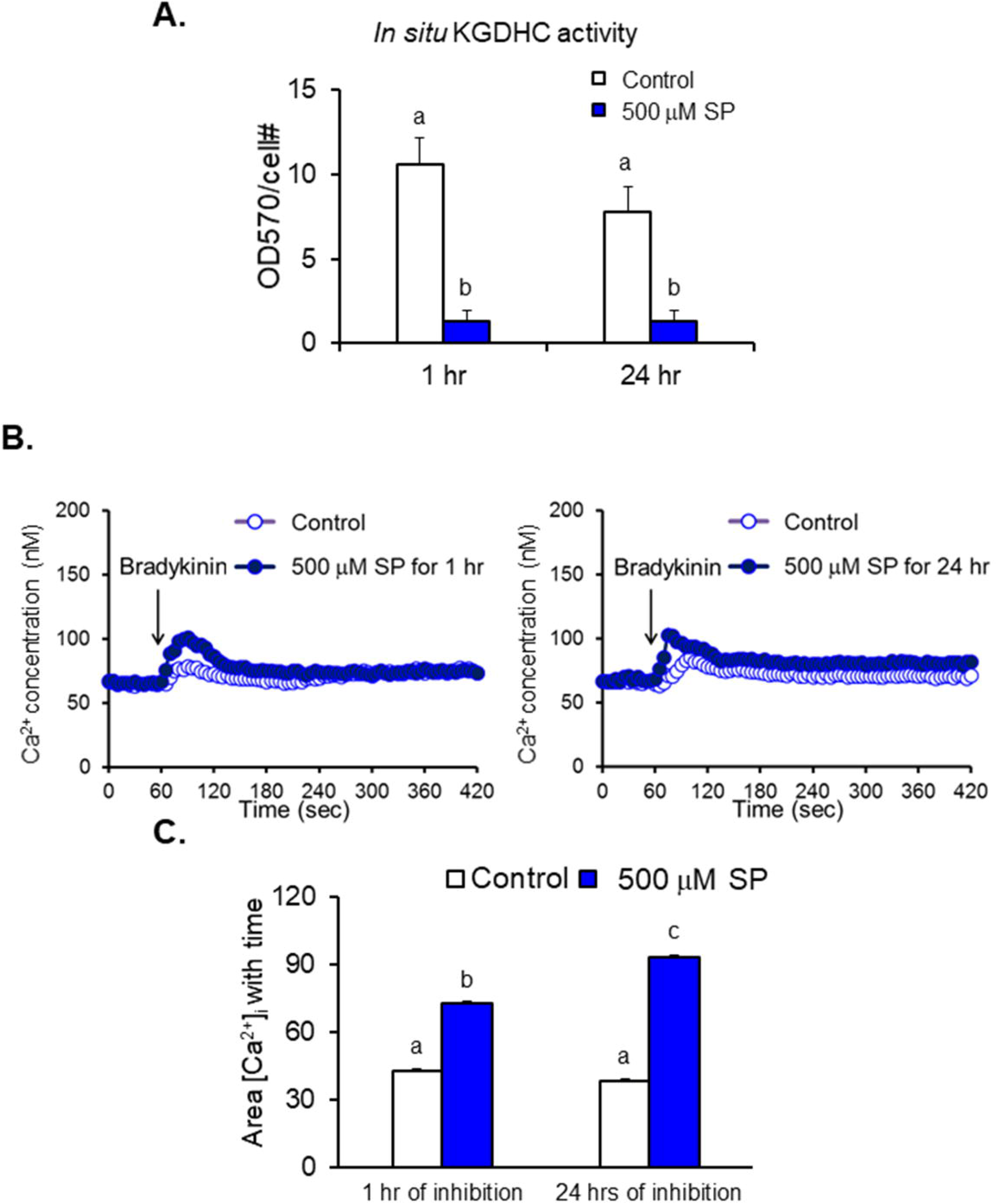
Reducing KGDHC exaggerated BRCS in neurons. (A) iPSC-derived neurons were cultured on 24-well plates at the seeding density of 5 × 10^4^ cell/well for four weeks. After four weeks of culture, KGDHC activity was measured by histochemistry (Panel A). Neurons were cultured on Delta T dishes for four weeks. After culture, the cells were pre-treated with SP (500 μM) for one hour (in BSS) or 24 hours (in completed media). After 1 hour of Fura-2 loading, the media were changed to calcium free BSS, and the calcium measurements were initiated and bradykinin (500 nM) after 1 min of basal [Ca^2+^]_i_ measurement (Panel B). Values are means ± SEM (n=627 and 603 cells for one hour and 24 hours of treatment, respectively). Different letters indicate values vary significantly (p<0.05) from the other groups by ANOVA followed by Student Newman Keul’s test. (Panel C).

### Consequences of reducing PDHC on BRCS in iPSC-derived neurons

iPSC-Derived neurons were treated with the specific PDHC inhibitor, dimethyl acetyl phosphonate (DMAP, 5 or 100 μM) for one hour (Park *et al*. 2000). First, we confirmed the inhibitory effect of DMAP with histochemistry. In these neurons, 5 μM or 100 μM DMAP inhibited PDHC activity by 75% and 76%, respectively (Figure 9A). 5 μM or 100 μM DMAP inhibited BRCS by 44.8% and 46.4%, respectively (Figure 9).

**Figure 9.**
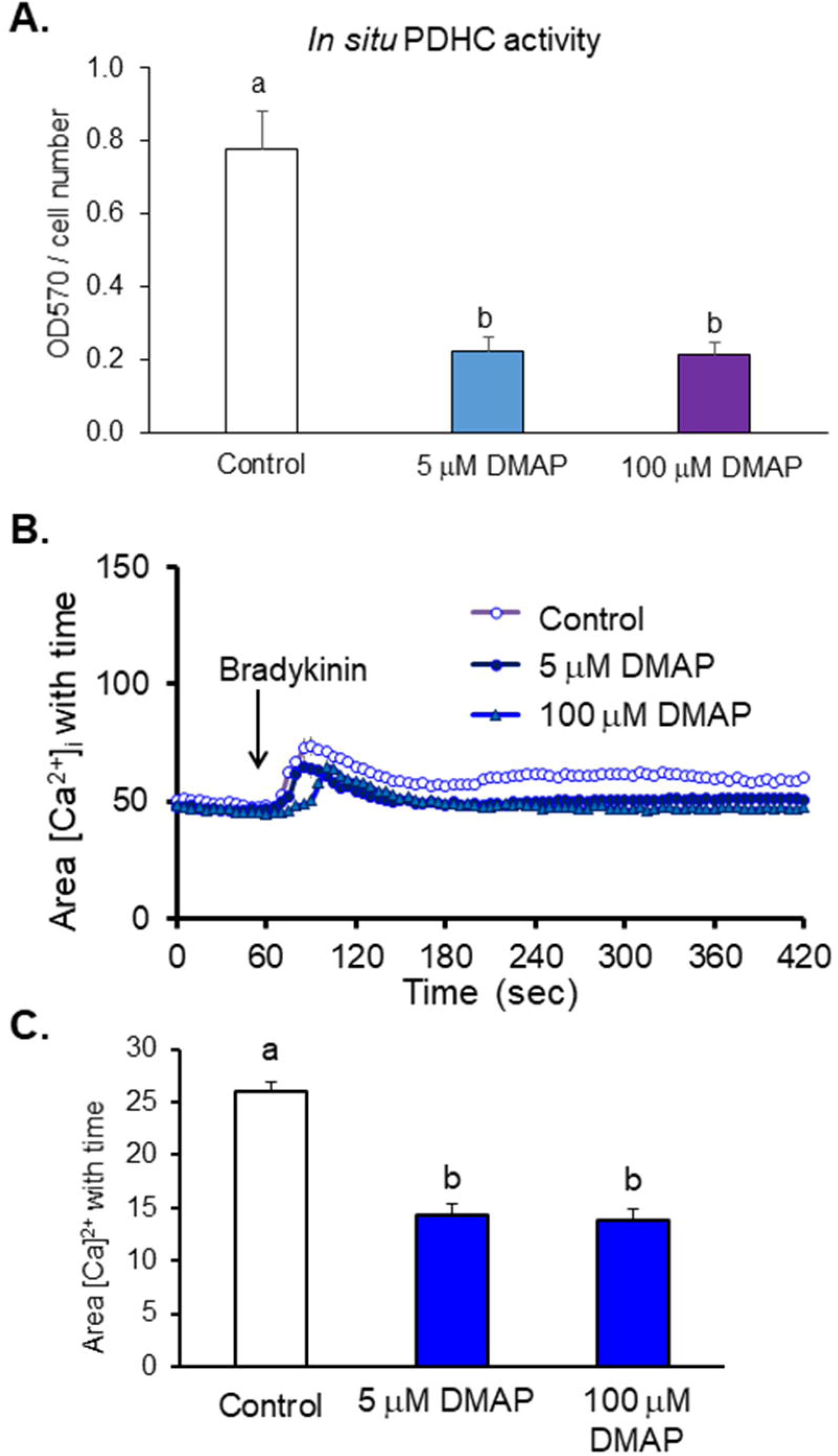
Reducing PDH diminished BRCS in iPSC derived neurons. iPSC-derived neurons were cultured on 24-well plates at the seeding density of 5 × 10^4^ cells/well for four weeks. After four weeks of culture, the activity of PDHC was measured by histochemistry (Panel A). Neurons were cultured on Delta T dishes for four weeks. After culture, the cells were pre-treated with DMAP (5, 100 μM) for one hour (in BSS). After 1 hour of Fura-2 loading, the media were changed to calcium free BSS, and the calcium measurements were initiated and bradykinin (500 nM) after 1 min of basal [Ca^2+^]_i_ measurement.(Panel B). Values are means ± SEM (n= 840 cells). Different letters indicate values vary significantly (p<0.05) from the other groups by ANOVA followed by Student Newman Keul’s test (Panel C).

### Consequences of reducing PDHC or KGDHC on resting cytosolic free calcium

The BRCS change in iPSC-derived neurons did not appear to be related to alter resting cytosolic free calcium, since changes in cytosolic free calcium were minimal when reducing KGDHC or PDHC in the absence IP_3_ agonists (Figure 10).

**Figure 10.**
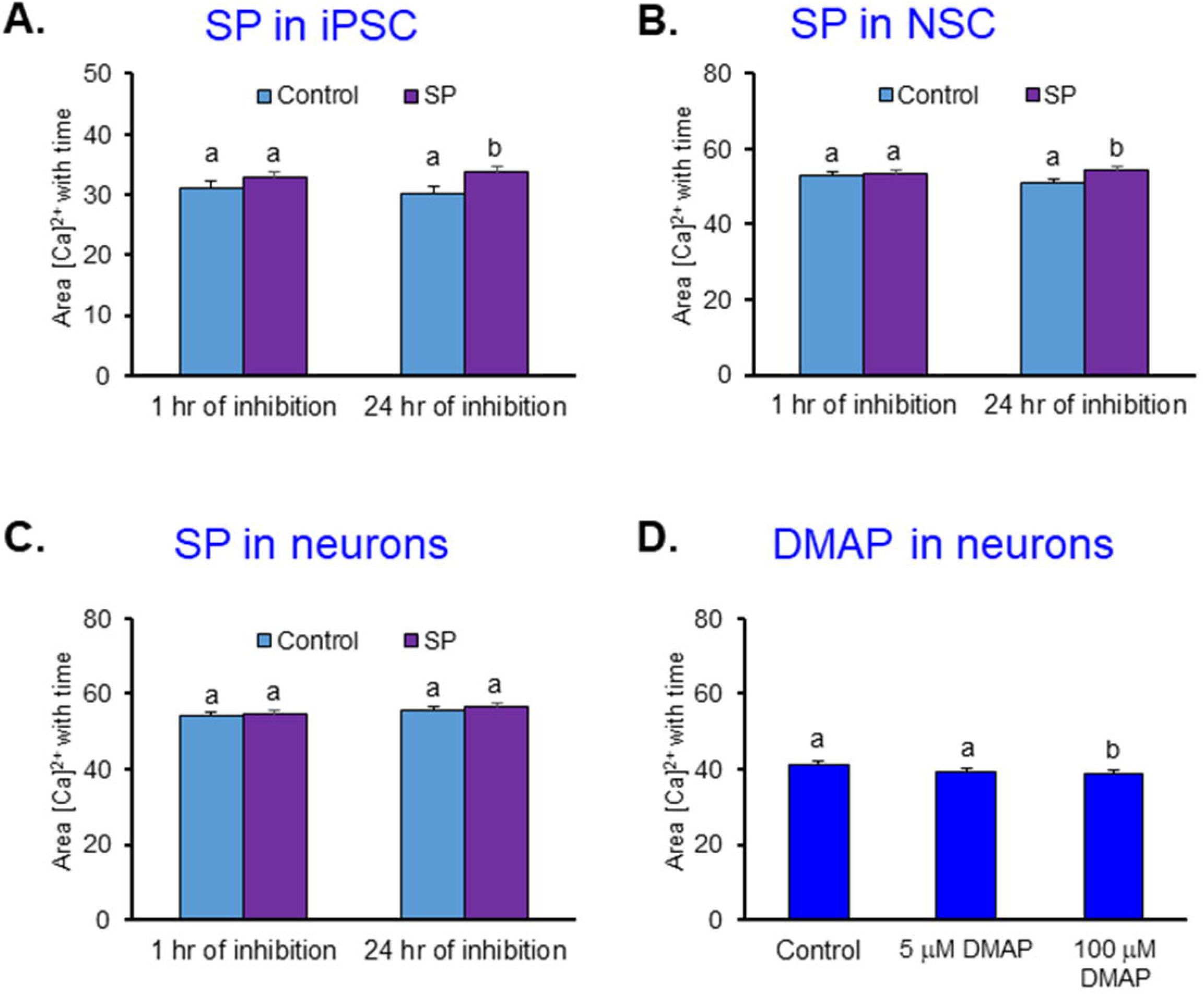
Reducing KGDHC or PDHC on basal calcium in iPSC, NSC, and iPSC derived neurons. Basal [Ca^2+^]_i_ in iPSC (A), NSC (B), and iPSC derived neurons (C)(D) were calculated from the same experiments in Figure 6–9. Values are means ± SEM. Different letters indicate values vary significantly (p<0.05) from the other groups by ANOVA followed by Student Newman Keul’s test.

## Discussion

These experiments are the first to report that the release of ER calcium stores and their relation to mitochondria differ in iPSC, NSC and neurons. Although the cells at all stages of development had IP_3_ releasable stores, the patterns were markedly different in NSC compared to iPSC and iPSC-derived neurons. Previous studies only showed the cytosolic calcium changes in iPSC-derived neurons for the study of AD (Prè *et al*. 2014; Duan *et al*. 2014), motor or dopaminergic neurons (Hartfield *et al*. 2014; Dafinca *et al*. 2016) or other neurodegenerative diseases (Hartfield *et al*. 2014; Schöndorf *et al*. 2014; Rabenstein *et al*. 2017; Prè *et al*. 2014). In addition to establishing differing patterns of BRCS at different stages of development, the experiments in this manuscript demonstrate that the mitochondrial link to BRCS varies. Reducing KGDHC in human iPSC-derived neurons exaggerated BRCS, but had no impact in NSC or iPSC. This difference may relate to the fact that mitochondria have a very different role in iPSC, neural stem cells and neurons. iPSC and neural stem cells primarily use glycolysis for energy, whereas neurons depend on mitochondria. BRCS are sensitive to select free radicals, and free radical production is much higher when cells use mitochondria for energy (Khacho *et al*. 2019; Intlekofer & Finley 2019). Thus, differences in ROS may underlie the differences in the sensitivity of the various cell types to inhibition of PDHC or KGDHC.

The cellular basis for the selective effects of inhibiting PDHC and KGDHC on BRCS in neurons is unknown. The selective effect makes it unlikely that this is a consequence of altered mitochondrial buffering of cytosolic calcium. Inhibiting PDHC will block acetyl-coA entry into the tricarboxylic acid cycle, which will deplete the substrates of the tricarboxylic acid cycle. On the other hand, blocking KGDHC may lead to accumulation of substrates such as alpha-ketoglutarate that leave the mitochondria and serve as signaling molecules throughout the cell. The current results in human neurons confirm our previous studies in rodent neurons that showed that inhibition of KGDHC exaggerates BRCS in an AD-like manner. Our previous studies in mouse cells showed that diminishing KGDHC increased BRCS under conditions that increased the mitochondrial NAD^+^/NADH ratio and elevated cytochrome c release (Gibson *et al*. 2012; Chen *et al*. 2017). The precise mechanism is unknown, but reductions in KGDHC diminish GTP (Kiss *et al*. 2013), and GTP is known to modulate ER-mitochondrial interactions (Hajnóczky *et al*. 1994). Furthermore, diminished KGDHC activity causes release of mitochondrial proteins (Banerjee *et al*. 2016) and post-translational modification of non-mitochondrial proteins (Yang *et al*. 2019). No other studies have examined the consequences of inhibiting PDHC on BRCS.

The relation of the increased BRCS with KGDHC inhibition and the elevation in BRCS in cells from AD patients and models of AD is not clear. The exaggerated BRCS with AD is well documented, and the current experiments further document a connection to KGDHC activity. Since KGDHC activity declines in AD brain, the decline may underlie the BRCS change in AD. The reduced KGDHC activity may also further exaggerate the increase in calcium in the AD cells. The exaggerated release of calcium from ER by activation of either ryanodine receptors (Stutzmann *et al*. 2004; Stutzmann *et al*. 2006) (Flucher *et al*. 1993) or IP_3_ receptors (Ito *et al*. 1994) (Chakroborty & Stutzmann 2014; Cheung *et al*. 2008; Del Prete *et al*. 2014) (Popugaeva & Bezprozvanny 2013) (Shilling *et al*. 2014) is among the most reproducible abnormalities that support a critical role of calcium in AD (Peterson C 1985). FAD mutant PS1 (M146L) and PS2 (N141I) interact with the IP_3_R Ca^2+^ release channel and exert profound stimulatory effects on its gating activity in response to saturating and sub-optimal levels of IP_3_ (Cheung *et al*. 2008). In addition, the membrane hyperpolarization associated with ER-calcium release is enhanced in mutant PS1 mice and reduces spiking activity and responsivity to synaptic inputs (Stutzmann *et al*. 2006; Oddo *et al*. 2003; Stutzmann 2007). The increased calcium leads to premature and severe defects in synaptic plasticity, behavior and cognitive function (Lacampagne *et al*. 2017). The calcium dysregulations seen in AD are not merely accelerated or amplified signaling changes, inevitable in old age, but rather, are novel and pathogenic changes to fundamental calcium signaling patterns (Stutzmann 2007). Reductions in KGDHC activity similar to those in AD would exaggerate these calcium effects in AD.

The lack of understanding of the molecular basis for the exaggerated BRCS in AD tissues makes it difficult to describe a molecular mechanism linking KGDHC and PDHC to the exaggerated BRCS. Our previous studies show that one possibility is the excess oxidants in AD cells. Select oxidants such as *t*-BHP (*tert*-butyl-hydroperoxide) or SNAP (S-nitroso-*N*-acetylpenicillamine) can cause AD-like changes in cytosolic calcium concentrations. AD fibroblasts have higher ROS production and BRCS (with *t*-BHP) than aged controls (Huang *et al*. 2010). Oxidant induced modification of protein thiol groups may underlie the AD-related exaggeration of the BRCS. These oxidants can alter BRCS by activating RyR (ryanodine receptor) transnitrosation of thiol containing proteins (Huang *et al*. 2005; Arnelle & Stamler 1995). Consistent with this idea are recent studies which show post-translational modifications of the ryanodine receptor by phosphorylation, oxidation and nitrosylation in brains of AD patients, and in two murine models of AD (Lacampagne *et al*. 2017). Whether this is also true of the IP_3_ receptor is not known. An increased number of ryanodine receptors appears important in mice with mutated presenilin; increased ROS expression in neurons contributes to increased calcium release, synaptic decline and increased RyR2 expression seen in the hippocampus and cortex. This increased IP_3_ receptor-evoked calcium release may involve the increase of stromal interaction molecule 1 (STIM1) expression and the decrease of STIM2 expression in the soma. These STIMs translocate to plasma membrane–ER contact sites, where they bind to Orai1 channels and open a Ca^2+^ influx pathway to slowly replenish ER Ca^2+^ stores using extracellular Ca^2+^(Mustaly-Kalimi *et al*. 2018; Schrank *et al*. 2019).

The effects of increased cytosolic calcium following release from ER calcium stores on the mitochondria is not clear. Mitochondria serve as an acute calcium buffer, so the exaggerated BRCS likely increases mitochondrial calcium concentration. In heart, increased intracellular Ca^2+^ concentration results in excess uptake of Ca^2+^ by the mitochondria and increased respiratory rate by the TCA cycle. However, this also leads to mitochondrial calcium overload as well as metabolic dysfunction (Schrank *et al*. 2019). Prolonged increases of Ca^2+^ release open the mitochondrial permeability transition pore (PTP) to release apoptotic signaling molecules (e.g., cytochrome C) (Schrank *et al*. 2019; Mustaly-Kalimi *et al*. 2018).

These results demonstrate that the relation of ER calcium to mitochondria changes with development. The results provide evidence in iPSC-derived human neurons that diminishing the activity of KGDHC causes AD-like changes in release of calcium from the endoplasmic reticulum.

## Methods

### Culture of iPSC, NSC and iPSC-derived neurons

We selected iPSC that were generated from fibroblasts (AG6842) obtained from the Coriell Institute (Camden, NJ). The fibroblasts were collected from a 75 year-old male donor, who was from a family bearing A246E PS1 mutation, but he did not have the mutation and was APO ε3/ε3. Fibroblasts were reprogrammed using four high-titer retroviral constructs prepared by the Harvard Gene Therapy Core Facility that encoded human Oct4, KLF4, SOX2 and c-Myc, respectively (Sproul *et al*. 2014). The iPSC colonies were initially selected by morphology, passaged several times to remove transformed cells and expanded before characterization. After iPSCs were expanded to multi-well format, they were characterized using a variety of quality control assays including karyotype and fingerprinting for cell-line genetics, alkaline-phosphatase (AP) enzymatic activity for reprogramming process, immunostaining for pluripotency markers, qPCR for endogenous pluripotent markers and viral transgene silencing, as well as APOE and PSEN1 genotyping (Sproul *et al*. 2014).

iPSC were cultured on dishes and plates with pre-coating (one hour at 37 °C) of Geltrex matrix solution (1:100 dilution) and the cells were cultured with StemFlex Medium for one or two days (ThermoFisher Scientific, Grand Island NY). iPSC were harvested with StemPro Accutase (ThermoFisher Scientific) for one minute at 37 °C and then cultured with StemFlex medium containing ROCK Inhibitor, Stemolecule Thiazovivin (1 μM; Reprocell, Beltsville, MD) for the first 24 hours of culture. After 24 hours, the medium was replaced with StemFlex Medium without ROCK Inhibitor, and medium was then exchanged daily. iPSC were cryopreserved in Synth-a-Freeze cryopreservation medium (ThermoFisher Scientific) and experiments were only performed using iPSC from passage 30 to 35.

NSC were induced from iPSC with a seven-day incubation with PSC Neural Induction Medium (NIM, Neurobasal Medium and Neural Induction Supplement; ThermoFisher Scientific) on Geltrex pre-coated plates. Media was changed daily. After induction, the NSC were harvested with StemPro Accutase for 5 min at 37 °C and cultured with Neural Expansion Medium (NEM, Neurobasal Medium, Advanced DMDM/F-12 and Neural Induction Supplement; ThermoFisher Scientific) containing Rock inhibitor Y27632 (5 μM, Sigma) for the first 48 hours of culture. NSC were cryopreserved in Synth-a-Freeze cryopreservation medium and neural differentiation was only preformed using passage five NSC.

For neuronal differentiation, the dishes or plates were pre-coated with Poly-L-ornithine (100 μg/mL) at 37 °C for two hours and Laminin (3 μg/mL) at 37 °C overnight. iPSC-derived neurons were differentiated from NSC (passage 5) for 28 days with Neuronal Differentiation Medium (Neurobasal Medium, 1 × B-27 supplement with antioxidant, 1 × GlutaMAX supplement, 2 × CultureOne supplement, 200 μM ascorbic acid, 1 × Anti-Anti, 20 ng/mL BDNF, 20 ng/mL GDNF; ThermoFisher Scientific) containing Rock Inhibitor Y27632 (5 μM) for the first 48 hours of differentiation at the seeding density of 2.5 × 10^4^ cell/cm^2^. The media was exchanged partially by removing half of the medium and replacing it with Neuronal Differentiation Medium twice weekly. Cells were typically used for neuronal markers characterization and BRCS after four weeks in culture. Differentiation was done at least 15 times.

### Immunostaining of cell specific cell markers

iPSC, NSC and iPSC-derived neurons were cultured on Delta T dishes (Bioptechs, Butler, PA) at different seeding densities and culture times (iPSC and NSC, 2 *×* 10^5^ cell/dish for two days; iPSC-derived neurons, 1 × 10^5^ cell/dish for four weeks). After culture, the cells were fixed with Image-iT Fixative Solution (4% formaldehyde; ThermoFisher Scientific) at room temperature for 15 min. After fixation, the cells were rinsed with DPBS (ThermoFisher Scientific) and blocked with blocking buffer (0.1% Triton X-100, 1% BSA in DPBS) at room temperature for one hour. After blocking, the cells were incubated with primary antibodies in blocking buffer at 4 °C overnight. The next day, the cells were rinsed with DPBS and incubated with secondary antibodies in blocking buffer at room temperature for one hour, following by staining with DAPI solution (ThermoFisher Scientific) at room temperature for 15 min. After DAPI staining, the dishes were mounted with ProLong Gold Antifade Mountant (ThermoFisher Scientific). The images were taken using a 20× objective under a Nikon 80i epifluorescence microscope. The table shows the details of primary and secondary antibodies.

| Primary Antibody | Vendor | RRID | Cat # | Dilution/dose |
| --- | --- | --- | --- | --- |
| OCT4 | ThermoFisher Scientific | AB_2534182 | A13998 | 1:400 |
| SSEA4 | ThermoFisher Scientific | AB_2533506 | 414000 | 10 µg/mL |
| Nestin | BD Biosciences | AB_396354 | 611658 | 1:1000 |
| SOX2 | ThermoFisher Scientific | AB_2533841 | 8-1400 | 3 µg/mL |
| SOX1 | R&D systems | AB_2239897 | AF3369 | 1:100 |
| PAX6 | ThermoFisher Scientific | AB_2533534 | 42-6600 | 1:200 |
| MAP2 | ThermoFisher | AB_2533001 | 13-1500 | 2 µg/mL |
|  | Scientific |  |  |  |
| Synaptophysin | EMD Millipore | AB_94786 | MAB329 | 1:500 |
| Neurofilament | ThermoFisher Scientific | AB_10984147 | MA5-14981 | 1:100 |
| **Secondary Antibody** | **Vendor** | **RRID** | **Cat #** | **Dilution/dose** |
| Alexa Fluor 488 donkey anti-mouse | ThermoFisher Scientific | AB_141607 | A21202 | 1:2000 |
| Alexa Fluor 488 donkey anti-goat | ThermoFisher Scientific | AB_2534102 | A11055 | 1:2000 |
| Alexa Fluor 594 donkey anti-rabbit | ThermoFisher Scientific | AB_141637 | A21207 | 1:2000 |
| Alexa Fluor 594 donkey anti-goat | ThermoFisher Scientific | AB_2534105 | A11058 | 1:2000 |

### Histochemical assay of KGDHC and PDHC

To estimate relative KGDHC and PDHC activity in intact cells, histochemical assays of these two enzymes were performed based on our pervious study (Park *et al*. 2000). iPSC, NSC and iPSC-derived neurons were seeded on 24-well plates at a seeding density of 5 × 10^4^ cells/well for different culture times (iPSC and NSC for 2 days, and iPSC-derived neurons for 28 days) in 0.5 mL of growth medium (details in culture section). On the day of each experiment, the medium was aspirated, and the cells were washed once with balanced salt solution (BSS) [140 mM NaCl, 5 mM KCl, 2.5 mM CaCl_2_, 1 mM MgCl_2_, 5 mM glucose and 10 mM HEPES (pH 7.4)]. The cells were then treated with SP (trisodium succinyl phosphonate, 100, 200 and 500 μM) or DMAP (5 or 100 μM) in 0.5 mL of BSS for 60 min at 37 °C. At the end of the treatment, the buffer was aspirated and the well was washed with 200 μL of Hank’s Balanced Salt Solution (HBSS) containing 0.05% (v/v) Triton X-100. Wells were incubated with 200 μL of either complete assay mixture or assay mixture without substrates as a negative control. Samples were incubated with the assay mixture for 40 min. After incubation, the treatment medium was aspirated and the cells were washed with Ca^2+^- and Mg^2+^-free HBSS. The dark blue formazan product was solubilized with 10% (w/v) SDS in 0.01 N HCl overnight in a CO_2_ incubator at 37 °C. The absorbance was read at 570 nm with a Spectra Max 250 model plate reader (Molecular Devices, Sunnyvale, CA).

### Measurement of K^+^ depolarization

Cells were loaded with 2 μM fura-2 AM (ThermoFisher Scientific) in BSS for one hour at 37 °C and rinsed twice with BSS (Gibson *et al*. 2002). Then, 2 mL of BSS was added to each dish and [Ca^2+^] was monitored on the stage of an inverted Olympus IX70 microscope at 37 °C with a Delta Scan System (Photon Technology International, Edison, NJ). Excitation wavelengths were alternated between 350 and 378 nm (band pass 4 nm) and emission was monitored at 510 nm with a Hamamatsu C2400 SIT camera (Hamamatsu, Hamamatsu City, Japan) at 5-second intervals. Basal [Ca^2+^] was measured for one minute. 50 mM KCl (Sigma) was added to depolarize voltage gated calcium channels to allow entry of extracellular calcium, which increases cytosolic calcium. The signal was measured for another 5 minutes. Each value was the average of 32 images taken within 5 seconds. Standard images of fura-2 solutions with minimum and maximum [Ca^2+^] were taken at the end of each day’s experiment to calculate the intracellular calcium concentrations (Huang *et al*. 1991).

### Measurement of BRCS

Cells were loaded with 2 μM fura-2 AM in BSS for one hour at 37 °C and rinsed twice with Ca^2+^-free BSS (140 mM NaCl, 5 mM KCl, 1.5 mM MgCl_2_, 5 mM glucose, 10 mM HEPES, 0.1 mM CaCl_2_,1mM EGTA, pH 7.4) (Gibson et al., 2002). Then, 2 mL of Ca^2+^-free BSS was added to each dish and [Ca^2+^] was monitored with the Delta Scan System. Basal [Ca^2+^] was measured for one minute. Bradykinin (100, 200 and 500 nM; Sigma) was added to release calcium from internal calcium stores and the signal was measured for another 5 minutes. Each value was the average of 32 images taken within 5 seconds. Standard images of fura-2 solutions with minimum and maximum [Ca^2+^] were taken at the end of each day’s experiment to calculate the intracellular calcium concentrations (Huang *et al*. 2010).

### Statistics

Microscoft EXCEL (RRID: SCR_016137) was used to prepare graphics and IBM SPSS Statistics 25 (RRID:SCR_002865) was used to test the statistical significance for one-way ANOVA with Student Newman Keul’s test.

## Abbreviations used

AP: alkaline-phosphatase
APOE4: epsilon 4 allele of Apolipoprotein E gene
AD: Alzheimer’s disease
*t*-BIIP: *tert*-butyl-hydroperoxide
BRCS: bombesin or bradykinin releasable calcium stores
BSS: balanced salt solution
CESP: carboxyethyl succinyl phosphonate
DMAP: dimethyl acetyl phosphonate
ER: endoplasmic reticulum
HBSS: Hank’s Balanced Salt Solution
IP_3_: inositol 1,4,5-trisphosphate
IP_3_R: inositol 1,4,5-trisphosphate receptors
iPSC: induced pluripotent stem cells
KGDHC: alpha-ketoglutarate dehydrogenase complex
NEM: neural expansion media
NIM: neural induction media
NSC: neural stem cells
PDHC: pyruvate dehydrogenase complex
PS-1: presenilin-1
PTP: permeability transition pore
RRID: research resource identifier (see scicrunch.org)
RyR: ryanodine receptors
SNAP: S-nitroso-N-acetylpenicillamine
SP: trisodium succinyl phosphonate
STIM: stromal interaction molecule
TCA: tricarboxylic acid cycle

## Acklowledgements and conflicts

This research was funded by NIH 2P01AG014930-15A1 and the Burke Foundation. None of the authors have a conflict of interest.

